# Ultrastructure Expansion Microscopy reveals the nanoscale cellular architecture of budding and fission yeast

**DOI:** 10.1101/2022.05.16.492060

**Authors:** Kerstin Hinterndorfer, Marine. H. Laporte, Felix Mikus, Lucas Tafur Petrozzi, Clélia Bourgoint, Manoel Prouteau, Gautam Dey, Robbie Loewith, Paul Guichard, Virginie Hamel

## Abstract

The budding yeast *Saccharomyces cerevisiae* and the fission yeast *Schizosaccharomyces pombe* have served as invaluable model organisms to study various fundamental and highly conserved cellular processes. While super-resolution (SR) microscopy has in recent years paved the way to a better understanding of the spatial organization of molecules in cells, its wide use in yeast models has remained limited due to the specific know-how and specialized instrumentation required, contrasted with the relative ease of endogenous tagging and live cell fluorescence microscopy in these systems. To facilitate SR microscopy in yeasts, we have extended the ultrastructure expansion microscopy (U-ExM) method to both *S. cerevisiae* and *S. pombe*, enabling 4-fold isotropic expansion in both systems. We demonstrate here that U-ExM allows the nanoscale imaging of the microtubule cytoskeleton and its associated spindle pole body (SPB), notably unveiling a conserved Sfi1p/Cdc31p spatial organization on the appendage bridge structure. In *S. pombe*, we validate the method by quantifying the homeostatic regulation of nuclear pore complex (NPC) number through the cell cycle. Combined with pan-labelling (NHS ester), which provides a global cellular context, U-ExM unveils the subcellular organization of the eukaryote yeast models *S. cerevisiae* and *S. pombe*. This easy-to-implement imaging with conventional microscopes provides nanoscale resolution and adds a powerful new method to the already extensive yeast toolbox.

## Introduction

*Saccharomyces cerevisiae* and *Schizosaccharomyces pombe* are unicellular ascomycete fungi on the order of 5-10 μm in size (Zakhartsev and Reuss, 2018) that share many fundamental features of cellular organization with animals. Decades of work using these model organisms has tremendously advanced our knowledge of fundamental cellular processes, including the regulation of the cell cycle, cell growth, DNA replication and repair, membrane trafficking, polarity, and signaling, enabled in large part by powerful genetic and molecular tools (Botstein et al., 1997; Hayles and Nurse, 2018). However, the resolution limits of classical live cell and immunofluorescence microscopy, combined with the small size of budding and fission yeast cells, have limited the full potential of cell biology studies in these models. The emergence of super-resolution microscopy has made it possible to circumvent this limitation, with a notable example being the use of Structured Illumination Microscopy (SIM) at 120 nm resolution to study the spindle pole body (SPB), a conserved fungal organelle reminiscent of the mammalian centrosome (Bestul et al., 2017; Burns et al., 2015; Unruh et al., 2018). Moreover, Stochastic Optical Reconstruction Microscopy (STORM) or Photo-Activated Localization Microscopy (PALM) of intracellular structures has achieved almost 50 nm resolution in *S. cerevisiae* and *S. pombe* (Arasada et al., 2018; Hajj et al., 2014; Prouteau et al., 2017; Ries et al., 2012). Nonetheless, the use of super-resolution microscopy to image yeast remains limited owing to complex image acquisition and processing routines and extensive, costly hardware requirements.

The recently developed expansion microscopy (ExM) method enables superresolution imaging using diffraction-limited microscopes. This protocol, including various extensions of the original, rely on the isotropic physical expansion of the biological sample (Chen et al., 2015; Gao et al., 2019; Ku et al., 2016). Among these protocols, Ultrastructure Expansion Microscopy (U-ExM) has been shown to preserve near-native cellular architecture (Gambarotto et al., 2019; Gambarotto et al., 2021; Zwettler et al., 2020). We reasoned that physical expansion of the yeasts *S. cerevisiae* and *S. pombe* by U-ExM could provide an easy and accessible super-resolution imaging method, much as it has for the microscopic parasites *Plasmodium, Toxoplasma* or *Trypanosoma* (Amodeo et al., 2021; Bertiaux et al., 2021; Tosetti et al., 2020). However, the presence of a robust cell wall in these organisms impedes proper or complete expansion, as shown for some bacterial strains or the nematode *C. elegans* (Lim et al., 2019; Yu et al., 2020). Recent ExM applications to fungi such as *Ustilago maydis* and *Aspergillus fumigatus* (Götz et al., 2020) as well as *S. cerevisiae* (Chen et al., 2021) pave the way for using expansion microscopy in these model organisms. However, such approaches include a full enzymatic treatment that digests the proteome, and more importantly, pre-expansion labeling which retains the linkage error (Zwettler et al., 2020). Post-labeling ExM protocol on as *S. Cerevisiae* has recently been reported (Korovesi et al., 2022); however, the resolution power of this method on different organelles, its compatibility with antibodies, and its feasibility in other yeast models remain to be assessed. Here, we develop a robust U-ExM protocol for budding and fission yeast, keeping the proteome intact and labelling post-expansion to reduce linkage errors (Hamel and Guichard, 2021). In addition, we combine U-ExM with NHS-ester pan-staining (M’Saad and Bewersdorf, 2020) to unveil the global cellular context for these two yeast models. We show that both *S. cerevisiae* and *S. pombe* can be expanded 4-fold, which we use to visualize the microtubule cytoskeleton at nanoscale resolution. We use U-ExM to probe the organization of the conserved Sfi1p•Cdc31p core module at the *S. cerevisiae* spindle pole body, and quantify nuclear pore complex (NPC) distributions at the nuclear surface as a function of cell cycle progression in *S. pombe.* Our results demonstrate that combining Pan-NHS-ester proteome staining with specific immunostaining provides an easy and accessible way to unveil the precise spatial distribution of intracellular structures.

## Results

### Establishing a U-ExM workflow for *S. cerevisiae* and *S. pombe*

In order to develop an easy protocol for expansion microscopy in the yeasts *S. cerevisiae* and *S. pombe*, we first optimized digestion of the cell wall (**Figure 1A**). Cells were grown in liquid culture to approximately OD of 0.4 and fixed with 3.7% paraformaldehyde (PFA) for 30 to 40 minutes at 21°C with shaking. After washes, the yeast cells were transferred into 1.2M sorbitol buffer prior to cell wall digestion in order to prevent osmotic shock and subsequent cell lysis. Digestion of the cell wall was performed by incubating the fixed cells with Zymolyase, an enzyme mix which digests the yeast cell wall (Kitamura and Yamamoto, 1972), with complete digestion confirmed by visual inspection. After washes in sorbitol buffer, the fixed and cell-wall digested cells were deposited on poly-D-lysine coated coverslips prior to further fixation with ice-cold methanol immediately followed by fixation in ice-cold acetone (**Figure 1A**; see material and methods section).

**Fig. 1.**
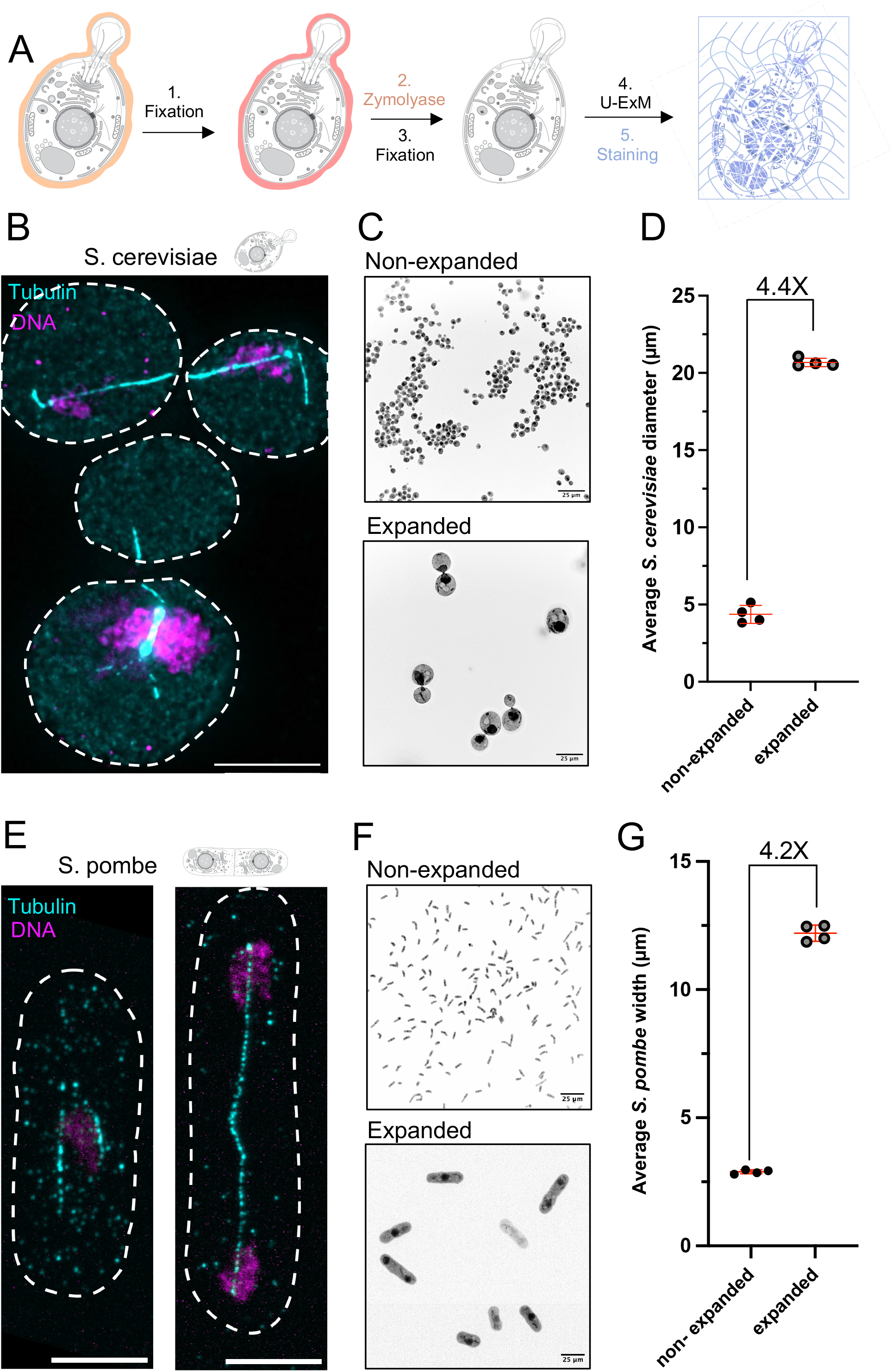
Workflow to perform U-ExM expansion microscopy in yeasts. (A) Schematic pipeline (swissBioPics) explaining the different steps of the U-ExM protocol on *S. cerevisiae* and *S. pombe*, including pre-fixation (1), cell wall digestion (2), further fixation (3), U-ExM (4) and staining (5). (B) Representative widefield images of expanded of *S. cerevisiae* cells stained with tubulin (cyan) and DAPI (magenta) displayed as maximum intensity projections. Scale bar: 10 μm. (C) Representative widefield images of *S. cerevisiae*, stained using the pan NHS-ester compound to visualize the entire cell, before (upper panel) and after expansion (lower panel). Scale bars: 25 μm. (D) Measurements of the average diameter of the entire *S. cerevisiae* cell before and after expansion, allowing the expansion factor calculation (n= 50-59 cells from 4 independent experiments; non-expanded= 4.37 +/− 0.58 μm and expanded= 20.68 +/− 0.27 μm). Note that the yeast cells are expanded 4-fold as expected. (E) Representative confocal images of *S. pombe* cells stained for tubulin (cyan), and Hoechst (magenta) displayed as maximum intensity projections. Scale bars indicate an actual distance of 10 μm. (F) Representative spinning disk confocal images of *S. pombe* cells stained using the pan NHS-ester compound to visualize the entire cell, before (upper panel) and after expansion (lower panel). Unadjusted length of scale bars: 25 μm. (G) Width measurements of *S. pombe* cells to determine the expansion factor. (n=72-98 cells from 4 independent experiment; non-expanded= 2.89 ± 0.08 μm and expanded= 12.2 ± 0.32 μm). Note that the yeast cells are expanded 4-fold as expected. Note that the scale bars in this figure indicate actual measured lengths; the following figures show lengths divided by the respective expansion factors.

Next, coverslips were briefly dried and directly placed into the anchoring buffer (FA/AA) for 5 hours at 37°C, followed by embedding into gels, denaturation and expansion ((Gambarotto et al., 2021) and **Figure 1A**). Gels were subsequently stained with YL1/2 tubulin antibody and DAPI to visualize microtubules and DNA respectively (**Figure 1B, E**). We found that the cytoplasmic microtubules emanating from the spindle pole body (SPB) as well as the mitotic spindle were nicely preserved under this condition in *Sc*, while in *Sp* better staining was achieved for mitotic spindles than for cytoplasmic microtubules, suggesting the need for additional optimization of the staining protocol (**Figure 1B, E**).

Next, to assess isotropic expansion and calculate the expansion factor, we measured the diameter of yeast cells before (4.37 μm +/− 0.58 and 2.89 μm +/− 0.08 μm for *Sc* and *Sp* respectively) and after expansion (20.68 μm +/− 0.27 and 12.21 +/− 0.3 μm for *Sc* and *Sp* respectively) using the NHS-ester compound that non-specifically labels the proteome, permitting straightforward recognition of cell boundaries (M’Saad and Bewersdorf, 2020). Importantly, we demonstrate that the yeast cells could be expanded ~4-fold as expected for this protocol (**Figure 1C-D, F-G**). We thus concluded that U-ExM allows a full expansion of both *S. cerevisiae* and *S. pombe* cell specimens.

### Visualizing features of the spindle pole body

The SPB is the microtubule-organizing center in fungi, which is functionally similar to the centrosome found in mammals (Ito and Bettencourt-Dias, 2018). Like its metazoan counterpart, the SPB is duplicated once per cell cycle and has microtubule nucleation abilities (Kilmartin, 2014). Our understanding of the *S. cerevisiae* SPB is grounded in electron microscopy (EM) (Kilmartin, 2014; O’Toole et al., 1999) and beautifully complemented with live imaging and more recently SIM (Burns et al., 2015; Geymonat et al., 2020; Unruh et al., 2018) as well as biochemical data (Rüthnick et al., 2021). The SPB is embedded in the nuclear membrane as a feature of yeast closed mitosis and is a cylindrical multi-layered organelle composed of outer, central and inner plaques (Jaspersen and Winey, 2004) (**Figure 2A, B**). Attached to one side of the central plaque of the SPB, and embedded in the nuclear envelope, is the half bridge or bridge depending on its length, with a key role during SPB duplication (Byers and Goetsch, 1975; Jaspersen and Winey, 2004) (**Figure 2B**). Decades of work in the yeast have uncovered most of the components of the SPB as well as their relative positions within this complex structure using either immuno-EM or GFP-tagging and fluorescence microscopy (reviewed in Helfant, 2002).

**Fig. 2.**
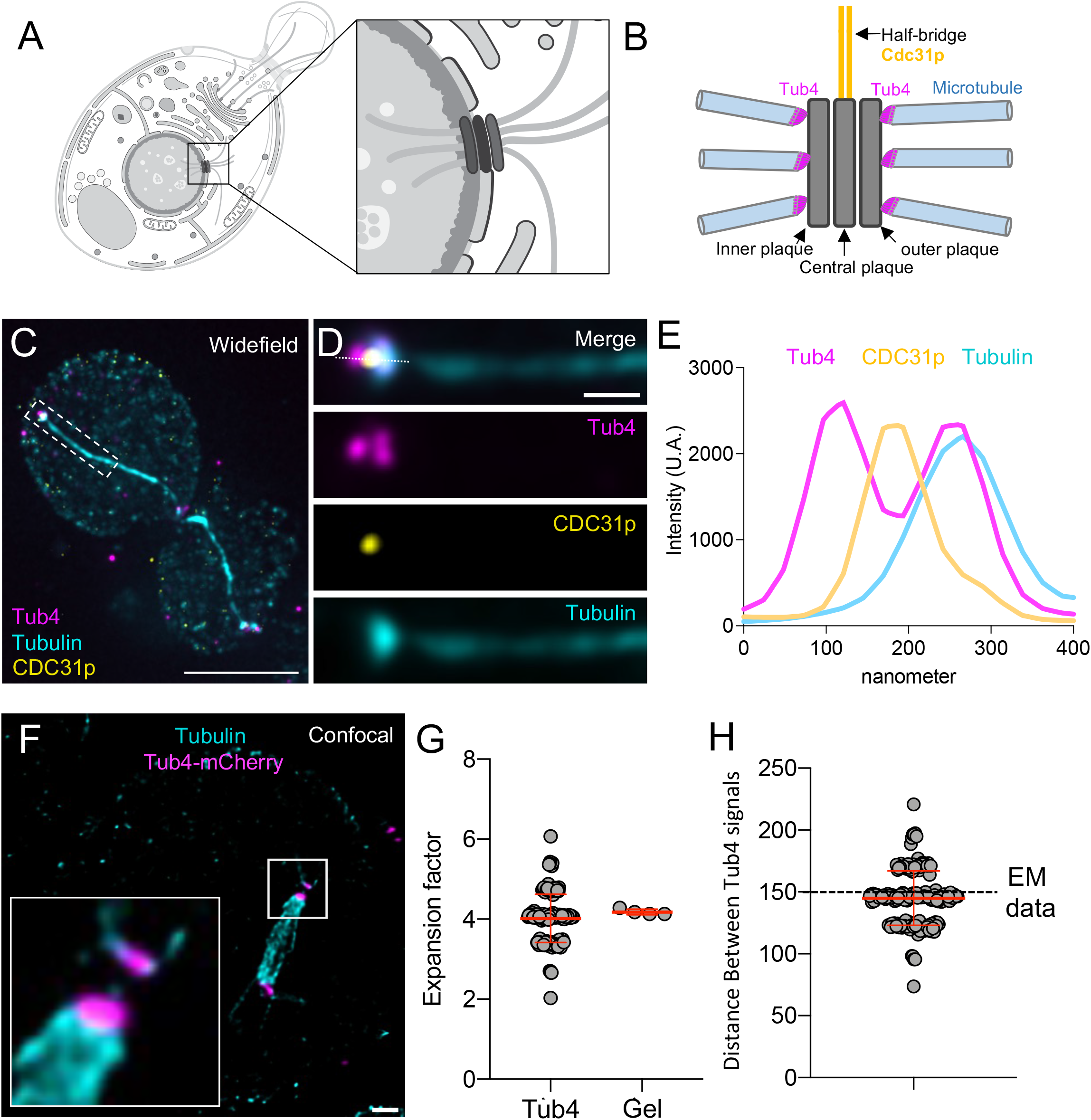
Visualizing the organization of the SPB in *S. cerevisiae.* (A) Schematic representation (swissBioPics) of a *S. cerevisiae* cell with an inset of its spindle pole body (SPB) embedded in the nuclear envelope. (B) Close up onto the SPB highlighting its unique spatial organization with the inner, central and outer plaques (grey) and its associated appendage structure called the half bridge, with its major component Cdc31p (yellow). Tub4 distribution, connecting the microtubules (cyan) to the outer plaque is indicated in magenta. (C) Representative widefield image of expanded *S. cerevisiae* cells stained for Tub4 (magenta), tubulin (cyan) and Cdc31p, as a marker of the half bridge structure (yellow). Scale bar: 2.5 μm. (D) Inset from the white dotted box in (C) highlighting the position of Tub4 (magenta), Cdc31p (yellow) relative to the microtubule (cyan) emanating from the SPB. Scale bar: 300 nm. (E) Plot profiles illustrating the distribution of the fluorescent signals for Tub4 (magenta), tubulin (light blue) and Cdc31p (yellow). Note that the signals can be easily distinguished, indicating that the two Tub4 distinct localization on the outer and inner plaques can be resolved using an epifluorescence microscope, as well as the half bridge structure seen using Cdc31p. (F) Confocal image of an expanded *S. cerevisiae* cell in mitosis, highlighting the mitotic spindle with its microtubules (cyan) and Tub4 (magenta). Scale bar: 300nm. The white box represents the inset, showing the easily distinguishable Tub4 signals. (G) Calculation of the expansion factor based on the measurement of an intracellular structure (left plot) or the diameter of the gel (right plot). For the measurement of the intracellular structure, the distance between Tub4 signals from post-expansion images was divided by the average distance between the two outer plaque from electron microscopy data (Byers and Goetsch, 1974). (n=179 cells from 4 independent experiments; Left: expansion factor= 4.02 +/− 0.604; Right: expansion factor=4.13 +/− 0.070. (H) Measurements of the distance between two Tub4 fluorescent signals (n=34-53 cells from 4 independent experiments; average calculated distance= 144.48 +/− 4.25).

To investigate the effective resolution we can achieve using U-ExM in yeast, we undertook an analysis of the g-tubulin homolog Tub4, an SPB component that is responsible for microtubule anchoring to the SPB (Spang et al., 1996). Tub4 is positioned on the inner and outer plaques of the SPB, at a measured distance of ~150 nm using SIM and electron microscopies (**Figure 2B**) (Burns et al., 2015; Byers and Goetsch, 1974). We wondered whether U-ExM would enable the resolution of the two plaques using conventional microscopes. *S. cerevisiae* cells expressing Tub4-mCherry were expanded and subsequently stained for tubulin to label microtubules and mCherry to position Tub4. We found that the two Tub4 fluorescent signals, representing the inner and outer plaques, could be easily distinguished using a widefield microscope (**Figure 2C-E**). Interestingly, we could also observe a distinct Cdc31p signal sandwiched between the Tub4 signals, indicating that we can resolve not only the inner and outer plaque of the SPB but also its adjacent half-bridge structure, of which Cdc31p is a component (**Figure 2C-E**) (Paoletti et al., 2003). Using confocal microscopy, we found that the two Tub4 plaques appeared even clearer with a measured distance of about ~144 nm +/− 5 after rescaling using the expansion factor, consistent with previous SIM and EM measurements (**Figure 2F, G**). Importantly, this finding also confirms that the expansion factor at the sub-cellular level is about 4-fold (**Figure 2H**). Altogether, our data demonstrate that U-ExM can achieve a similar or better resolution than what has been previously achieved using SIM.

### U-ExM resolves the Sfi1p/Cdc31p complex at the spindle pole body

In addition to Cdc31p, the sole centrin in budding yeast, the SPB half-bridge/bridge also contains Sfi1p (Bouhlel et al., 2015; Burns et al., 2015; Kilmartin, 2003; Li et al., 2006). This core module is critical for SPB duplication in yeast (Bouhlel et al., 2015; Kilmartin, 2003; Li et al., 2006; Rüthnick et al., 2021) while this function does not seem to be conserved in mammals (Bouhlel et al., 2021; Kodani et al., 2019). Immuno-EM analysis as well as SIM microscopy have localized the Sfi1p•Cdc31p complex in *S. cerevisiae* at the level of the bridge structure (Bestul et al., 2017; Li et al., 2006; Paoletti et al., 2003). Parallel bundles of elongated Sfi1p proteins (approximately ~60 nm long) form the core of the half bridge structure (Burns et al., 2015; Li et al., 2006; Rüthnick and Schiebel, 2018). The Sfi1p filaments are aligned with their N-termini facing the SPB and with their C-termini at the central region of the bridge (Seybold et al., 2015). This complex structure is stabilized and reinforced by the binding of the centrin Cdc31p along Sfi1p filaments through conserved binding sites (Kilmartin, 2003; Rüthnick et al., 2021; Seybold et al., 2015) (**Figure 3A**). During SPB duplication in anaphase, dephosphorylation of Sfi1p triggers antiparallel C-terminus to C-terminus dimerization and consequently, the elongation of the half bridge into the long bridge structure of ~ 120 nm (Kilmartin, 2003; Rüthnick et al., 2021). This unique organization of Sfi1p molecules orient all Sfi1p N-termini towards the SPBs, old and new, while all the Sfi1p C-termini face the center of the bridge (Kilmartin, 2014).

**Fig. 3.**
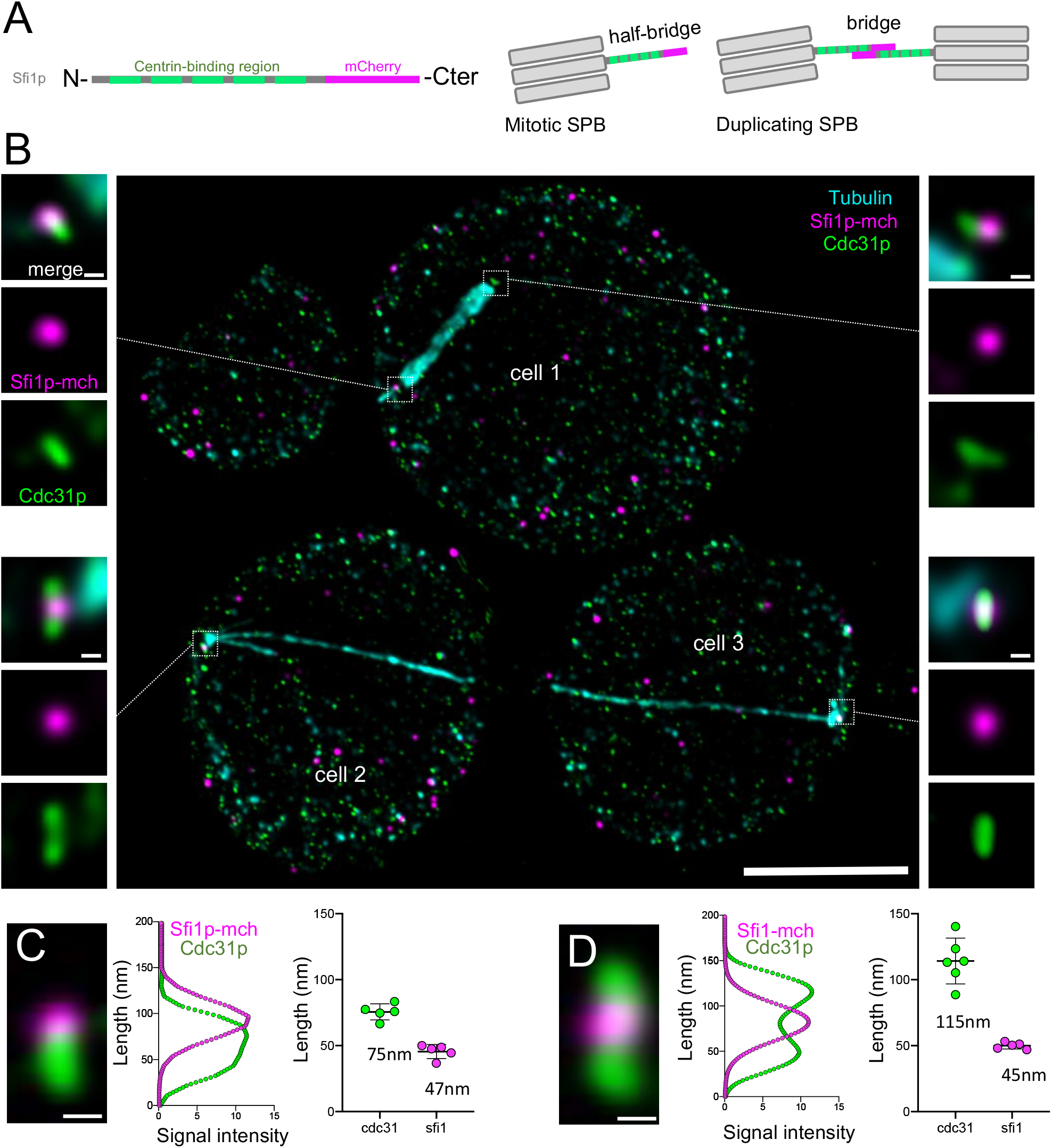
Resolving the conserved Sfi1p/Cdc31p complex at the SPB. (A) Schematic representation of the Sfi1p molecule, with the centrin-binding regions in green and the mCherry-Cterminal tag in magenta. In the half bridge structure, Cdc31p is close to the SPB while the C-termini of Sfi1p is external while in the full bridge, Sfi1p C-termini are placed in the center of the bridge. (B) Representative confocal image of expanded *S. cerevisiae* cells stained for tubulin (cyan), Sfi1p-mCherry (magenta) and Cdc31p (green). Insets are from the squared white boxes. Note that Sfi1p can be visualized as an external dot compared to the Cdc31p rod signal in the half bridge structure configuration (top insets), while it is found in the center of the Cdc31p extended rod signal in the full bridge (lower insets). Scale bars: 2.5 μm and 50 nm (insets). (C, D) Plot profiles and measurements of the length of the Sfi1p or Cdc31p signals in the half bridge (C) or full bridge (D) configurations. Scale bars: 50 nm.

We decided to use this Sfi1p•Cdc31p module as another test of the ability of U-ExM to increase resolution (**Figure 3A**). We engineered a Sfi1p-mcherry strain and stained expanded yeast for mcherry, to position Sfi1p C-termini, Cdc31p and tubulin, to mark the microtubules. We found that Sfi1p and Cdc31p positions can be readily distinguished either in the half-bridge or full-bridge configurations (**Figure 3B**). Confocal imaging of the expanded cells revealed a rod-shaped Cdc31p-positive structure either ~75 nm or ~115 nm long, which we hypothesize corresponds to the half-bridge or full-brige structure as predicted from the litterature (Kilmartin, 2003; Kilmartin, 2014; Li et al., 2006; Rüthnick et al., 2021) (**Figure 3C, D**). In contrast, the Sfi1p signal, which corresponds only to the location of its C-termini, was found either at the end of the Cdc31p signal (top panels) or in the center of the Cdc31p signal (bottom panels) with a constant length of ~45 nm, thus marking the extremity of the half-bridge pointing away from the SPB (**Figure 3A, C and D**). Altogether, we concluded that U-ExM applied to *S. cerevisiae* allows accurate imaging of SPB-associated appendage structures at nanoscale resolution.

### U-ExM enables the analysis of nuclear pore complexes throughout the cell cycle in *S. pombe*

The nuclear pore complex (NPC) is a deeply conserved, massive protein complex that regulates exchange between the nucleoplasm and cytoplasm, and is involved in determining various aspects of nuclear architecture (Hampoelz et al., 2019). Extensive EM studies have been carried out to characterize their composition, structure and, more recently, their dynamic nature in altered cellular states (Zimmerli et al., 2021). Due to their eight-fold symmetry and well-defined size, mammalian NPCs have been used as gold standards to validate super-resolution techniques such as PALM/STORM (Sabinina et al., 2021). In yeasts, however, their doughnut-like shape has so far been difficult to visualize using conventional microscopy approaches. NPCs largely regulate the nuclear transport of biomolecules and are involved in mitotic regulation by facilitating either a general nuclear envelope breakdown (NEBD) – in open mitosis – or a localized one in fission yeast closed mitosis (Dey et al., 2020). To maintain homeostatic NPC densities, the insertion of newly synthesized complexes is tightly regulated. Due to the small size of the yeast nucleus, this could only recently be quantified using 3D SIM (Varberg et al., 2022). We asked whether we could obtain matching results using U-ExM and conventional microscopy.

Staining of the NPC outer ring protein Nup107 in expanded *S. pombe* revealed individual NPCs, even when using widefield microscopy (**Figure S1**) and allowed us to easily quantify the number and diameter of NPCs throughout the cell cycle with confocal microscopy (**Fig. 4**). We measured the average size of individual pores to be 73.8 nm **(Figure S1)**, in line with the cryo-EM studies that provide the gold standard (Zimmerli et al., 2021). The NPC counts closely matched published measurements from 3D SIM (G1/S: 85 ± 10, Early G2: 92 ± 14, Mid G2: 120 ± 17, Late G2/M: 135 ± 21, Late M: 73 ± 14 NPCs, respectively). We expect that we are undercounting due to NPC clustering, an issue also faced by the 3D SIM analysis – one that could be addressed in the future by applying SIM or STED microscopy to expanded samples to further boost resolution.

**Fig. 4.**
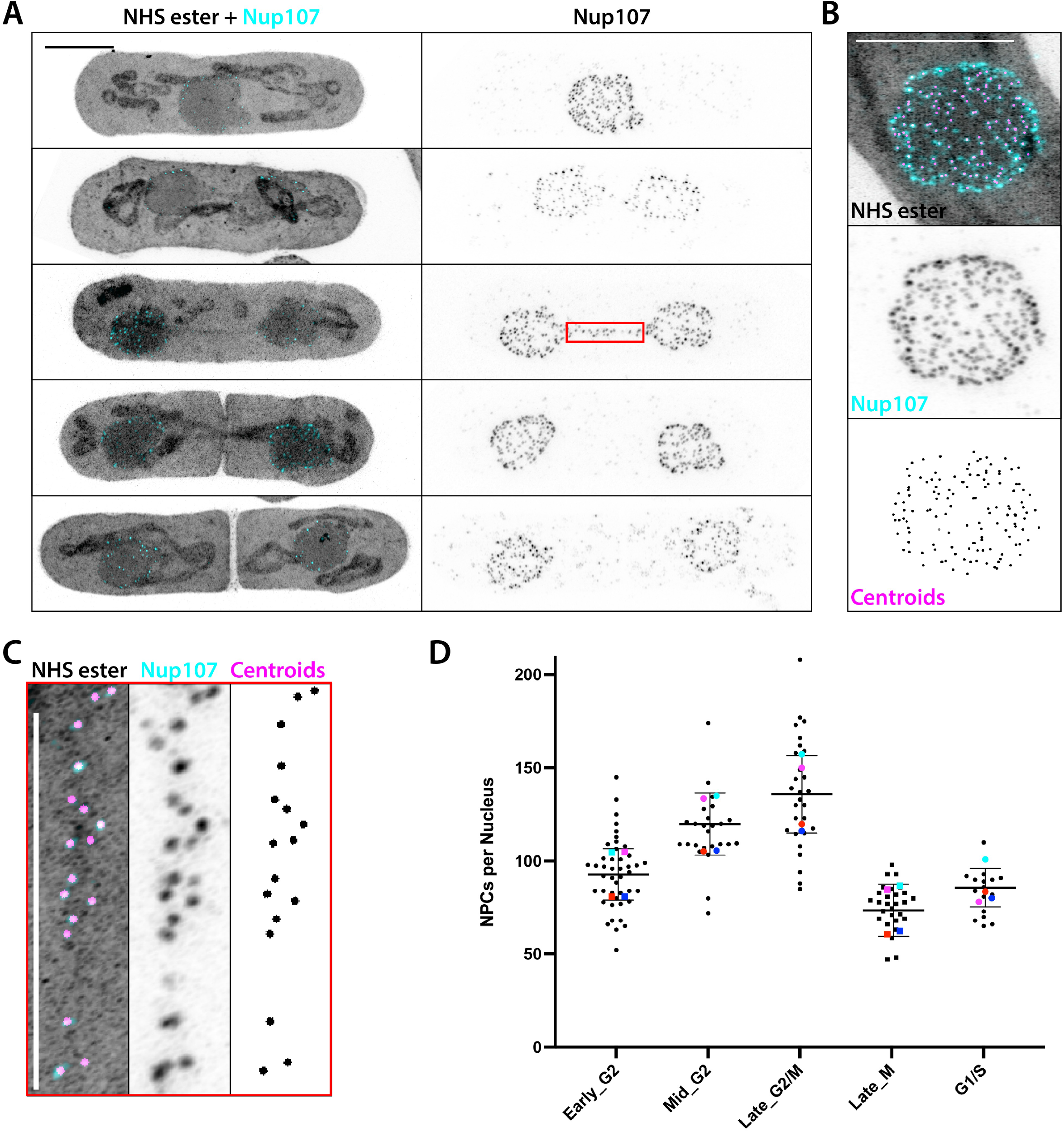
Quantification of fission yeast NPCs throughout the cell cycle. (A) Maximum intensity projections of the different stages in *S. pombe* division imaged at an AiryScan confocal system labelled with NHS ester (grey) and for Nup107 (cyan). The mitotic bridge can be distinguished in both channels (panel three and inset C), and pan-labelling easily differentiates fully closed septa. Scale bar: 2.5 μm. (B) Demonstration of the NPC segmentation displayed as a sum intensity projection of NHS ester (gray), Nup107 (cyan) and centroids from segmented Nup107 signals (magenta). Scale bar: 2.5 μm. (C) Zoom into the nuclear bridge from A showing individual NPCs and centroids derived from the analysis pipeline. Scale bar: 2.5 μm. (D) Quantification of total NPC numbers per nucleus in the different stages of the fission yeast cell cycle, coloured dots represent the averages from independent experiments. (n=35-53 cells from 4 independent experiment. G1/S: 85 ± 10, early G2: 92 ± 14, mid G2: 120 ± 17, late G2/M: 135 ± 21, late M: 73 ± 14 NPCs).

### Combining Pan NHS-ester staining with U-ExM to map the subcellular organization of budding and fission yeast

Finally, in combination with specific antibody or dye labeling, we carried out pan-NHS-ester staining, to map the global cellular context in expansion microscopy (Bertiaux et al., 2021; M’Saad and Bewersdorf, 2020; Mao et al., 2020), with U-ExM (**Figure 5**). We found that as in other model systems, NHS-ester staining unveils the general organization of the cell, and highlights specific structural elements of the cell such as the nucleus, the bud neck, and mitochondria (**Figure 5A**). By acquiring a 3D confocal stack of a dividing budding yeast, it was also possible to identify mitochondria traversing the bud neck exhibiting a local constriction (**Figure 5B**). However, the vacuole, a large membranous structure present in the budding yeast was not detectable using the pan-NHS ester staining, possibly due to its dilute composition or incomplete fixation (**Figure 5**). In *S. pombe*, the NHS ester preferentially stained the nucleoplasm and mitochondria (**Figure 4A, 5C**). Additionally, the SPB as well as the mitotic bridge, a thin connection formed by the nuclear envelope during mitosis, were clearly visible (**Figure 4A, 5C and Figure S1**). This additional information allowed us to easily determine if fluorescent signals were likely to derive from labelled nucleoporins, especially in mitotic cells where a subpopulation of NPCs is relocalised to the center of the mitotic bridge (Dey et al., 2020), a population that could be easily visualized and quantified using U-ExM.

**Fig. 5.**
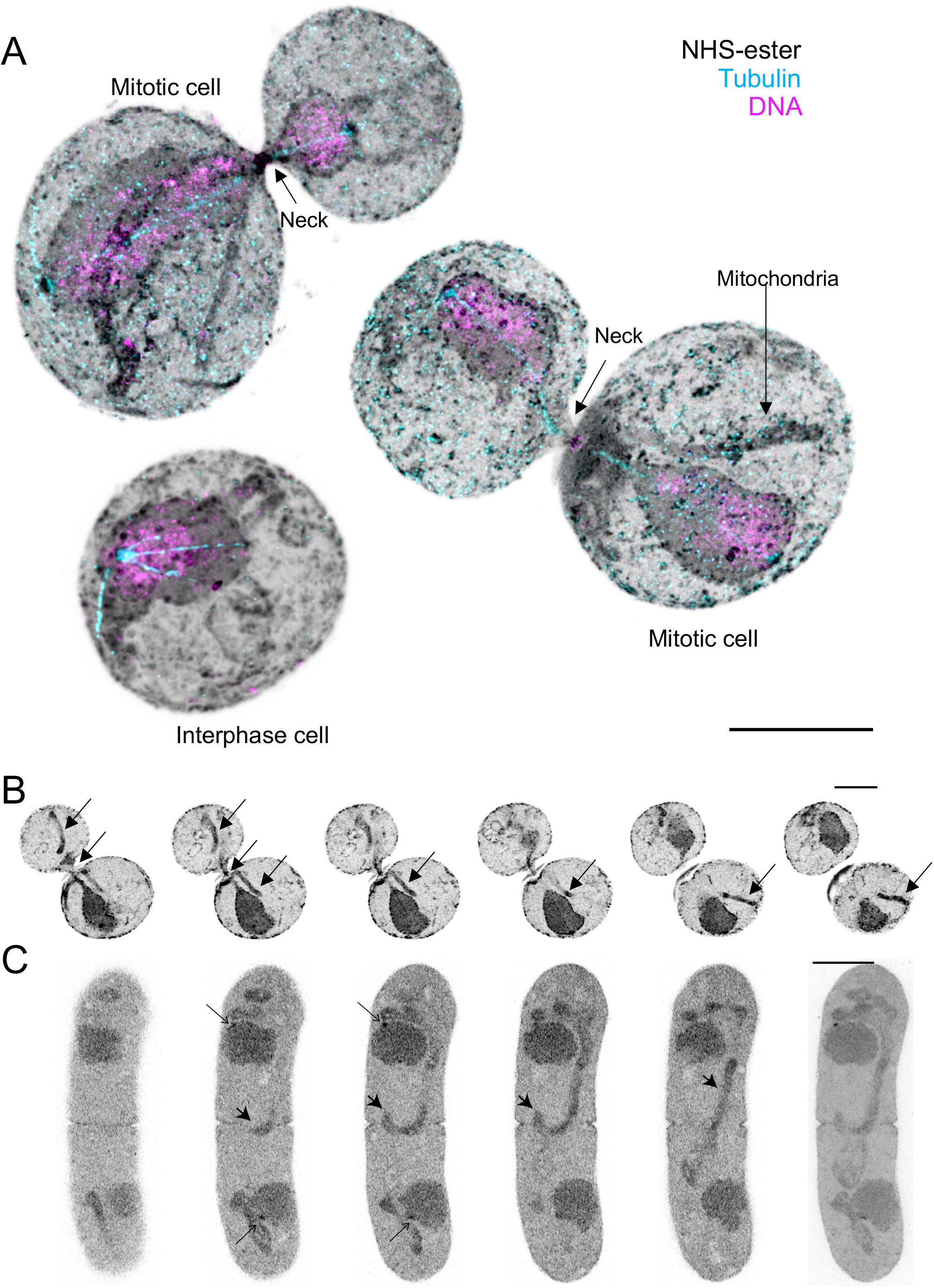
Pan NHS-ester and specific antibody labeling staining coupled to U-ExM unveils the spatial organization of the yeast. (A) Examples of expanded *S. cerevisiae* yeast cells stained with NHS-ester to provide a global cellular context, tubulin to mark microtubules (cyan) and DNA, acquired with a confocal microscope. Note that the cell contour can be easily recognizable as well as the bud and the neck separating the two cells. Elongated structures that we interpret as mitochondria can also be seen spanning the entire cell or even crossing the neck when the two cells are still connected at the end of cell division. Scale bar: 2.5 μm. (B) Montage of successive images of a confocal stack from *S. cerevisiae* (images are separated by 150 nm). The arrows indicate the position of a mitochondria that appears to pass through both cells and constricted at the level of the neck. (C) Montage of successive images of a confocal stack from a *S. pombe* cell in the process of forming the septum separating the two daughter cells (images are separated by 1.5 μm). Mitochondria (solid arrow) can still be found in the area of separation and the pan-labelling allows easy recognition of the SPBs at both daughter nuclei (open arrows). The last panel shows the maximum intensity projection of the entire stack with the scalebar indicating 2.5μm.

## Discussion

Here we present a straightforward approach to apply U-ExM to the budding yeast *S. cerevisiae* and the fission yeast *S. pombe.* Our results demonstrate that U-ExM can be successfully implemented following successive fixations and cell wall digestion steps prior to the application of the regular expansion protocol. With this method, we could visualize the nanoscale organization of the microtubule cytoskeleton. For example, we could observe distinct Tub4 fluorescent signals at each face of the spindle pole body, illustrating the power of U-ExM to resolve small structures. Another striking feature revealed by U-ExM is the conserved Sfi1p•Cdc31p complex, which displays a canonical organization at the bridge. Using U-ExM coupled to confocal imaging, we could resolve the position of Sfi1p C-termini at the center of the bridge as well as that of Cdc31p along the bridge. We could resolve nuclear pore complexes and determine their number throughout the cell cycle stages of fission yeast, demonstrating the possibilities U-ExM offers, in combination with conventional microscopy setups. Considering the ease of genetic manipulation in both yeast systems, one may also take advantage of these for expansion microscopy by using immunostaining directed against commonly used protein tags, such as mCherry, as demonstrated above. This will negate the need for protein-specific antibodies and further streamline the process of imaging expanded samples.

Nonetheless, it is worth highlighting a common limitation imposed by chemical fixation upon all imaging protocols, including U-ExM. Chemical fixation is a relatively slow process at the molecular level that can affect organelle morphology – for example, the vacuole, which is mainly composed of water (Frankl et al., 2015). Closer inspection of the overall nuclear envelope morphology in expanded fission yeast it might be affected by the current fixation method. Therefore, it would be very interesting in the future to adapt and implement the new method of cryo-fixation coupled to U-ExM, called Cryo-ExM (Laporte et al., 2022) to the yeast expansion protocol presented here. Indeed, cryo-fixation alleviates the need for specific chemical fixations and allows preservation of native cellular organization. The approach will consist of first using the well-established protocol of high pressure freezing (HPF) of the yeast cells, then processing of the sample for freeze substitution followed by U-ExM, as described previously (Laporte *et al,* 2022). In the future, coupling HPF and U-ExM might prove to be very powerful to image all types of molecular assemblies or sub-cellular structures in microbial model systems, including budding and fission yeast.

In summary, we propose that expansion microscopy applied to budding and fission yeast might become a useful tool to precisely study spatial organization of macromolecular complexes using conventional microscopes. This will undoubtedly help us understand basic mechanistic principles behind processes that are difficult to study owing to the small size of the yeasts.

## Acknowledgments

We thank Susanne Borgers and the Centriole lab for helpful discussions and technical advices as well as Vincent Louvel, Omaya Dudin, Hiral Shah and Jana Helsen for critical reading of the manuscript. FM and GD acknowledge the European Molecular Biology Laboratory (EMBL) for support, the EMBL Advanced Light Microscopy Facility (ALMF) for technical input. This work was supported by the ERC StG 715289 (ACCENT), the Swiss National Foundation (SNSF) 310030_205087 attributed to P.G. and VH and and (SNSF) 51NF40-185898 NCCR Chemical Biology – Visualisation and Control of Biological Processes Using Chemistry (phase III) to K.H. and R. L.

## Author contributions

K.H. performed all the *S. cerevisiae* experiments described in the paper under the supervision of M.H.L and L.T.P. and with support from M. P. and C. B.. FM performed all *S. pombe* experiments. G.D., V.H., P.G. and R.L. conceived, designed and supervised the project. All authors wrote and revised the final manuscript.

---

**Fig. S1.**
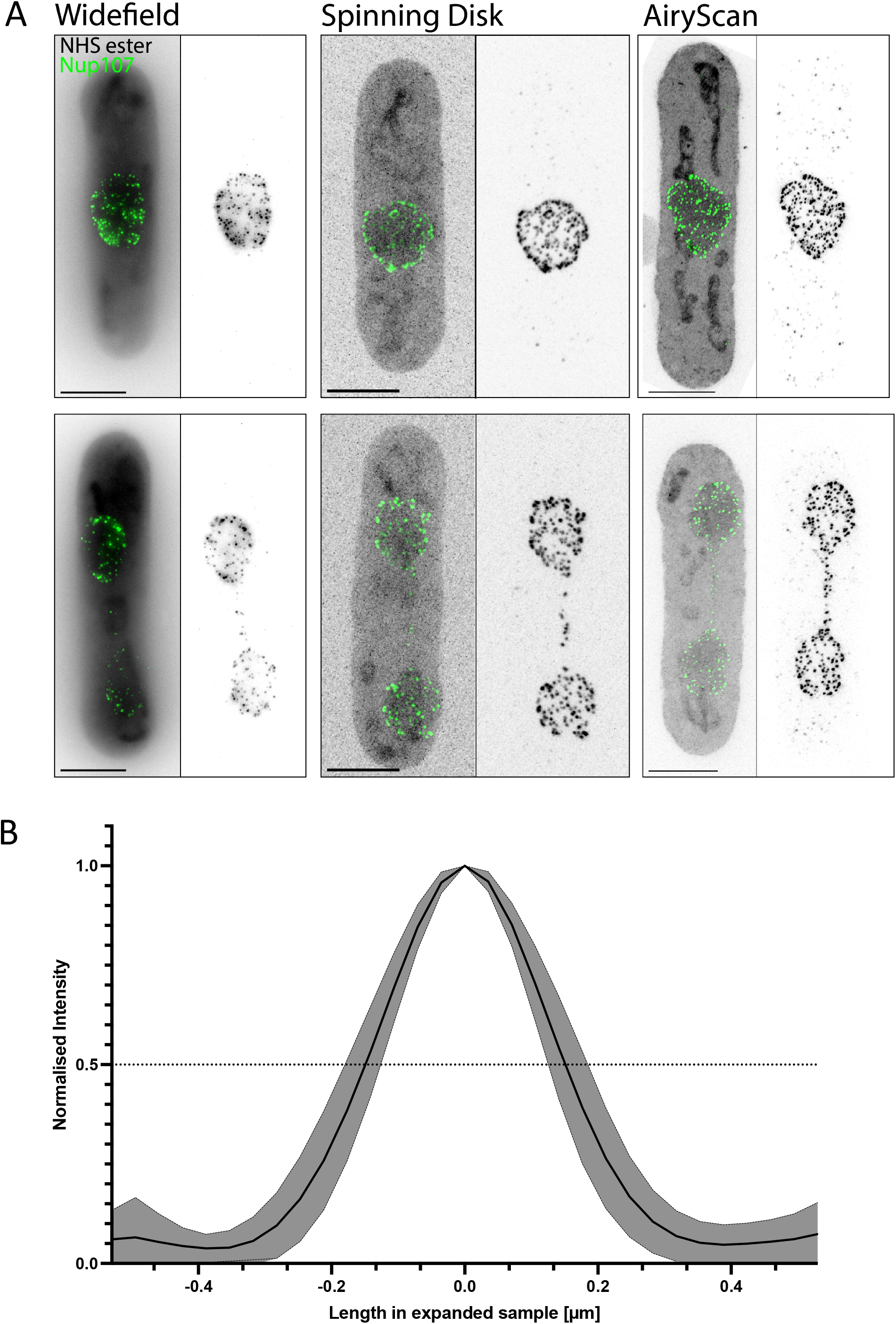
U-ExM allows reliable visualisation of small structures using different microscopy techniques: (A) Comparison of widefield (left), spinning disk confocal (middle), and AiryScan confocal (right) microscopy on representative *S. pombe* cells that are either in interphase or mid-mitosis. NPCs are labelled using a Nup107 antibody (green), NHS ester pan-labels the entirety of the cell (grey). Even widefield microscopy allows to distinguish signals we presume to be individual NPCs. Scale bars: 2.5 μm. (B) Plot of normalised fluorescence intensity derived from line profiles measured across Nup107 signals in polar nuclear planes (NPCs facing the detector) to determine the size. The width at half maximal intensity (0.5, dotted line) was determined to be 309.9 nm, which by dividing through the expansion factor of 4.2-fold indicates a signal size of 73.8 nm (n=129 NPCs).

## Materials and Methods

### Plasmids and primers

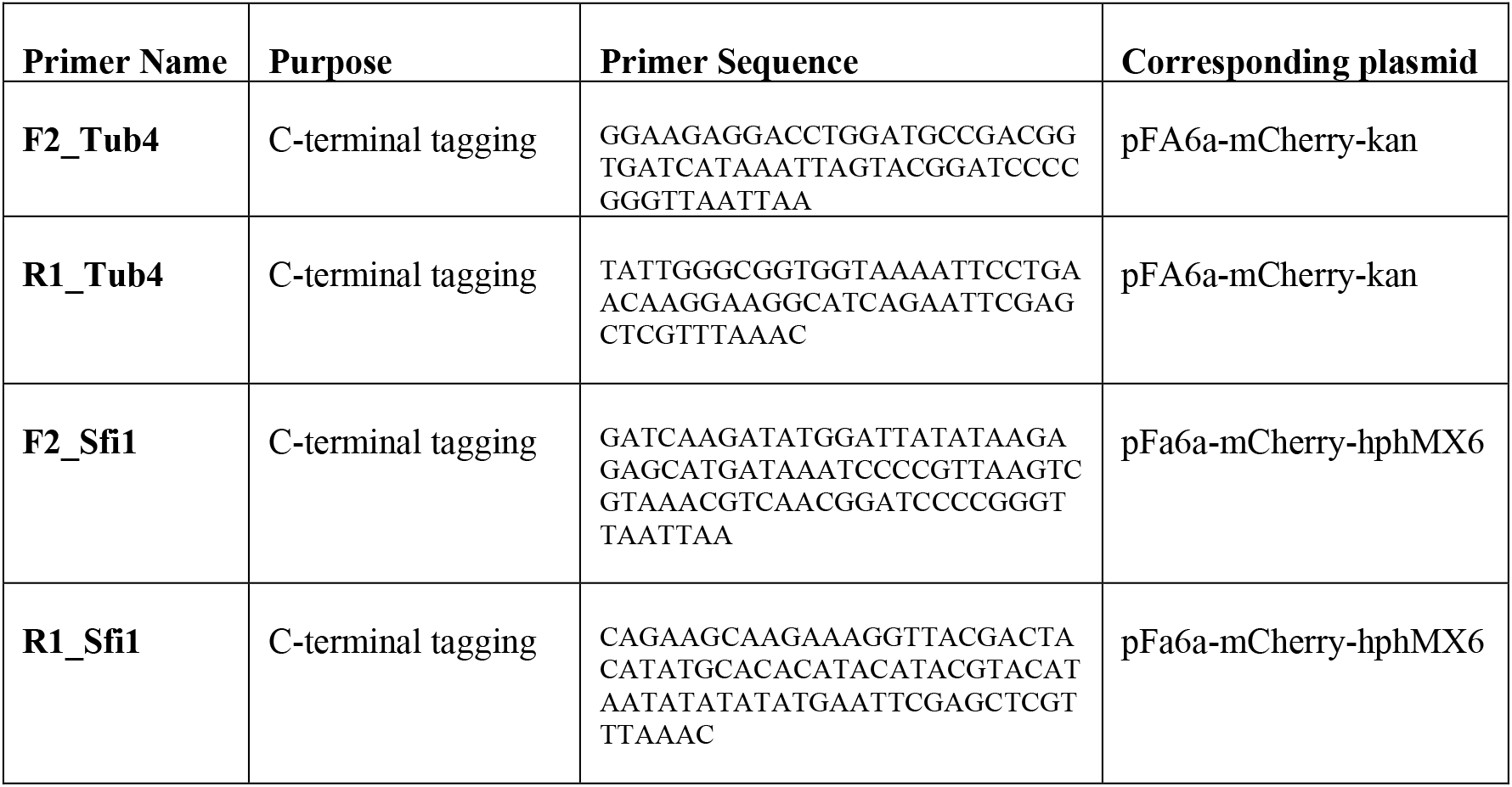

### Yeast strains and culture

*Schizosaccharomyces pombe* 972h-

The following strains were engineered: Sfi1-mcherry: TB50 Sfi1-mcherry::(HPH) Tub4-mcherry: TB50a Tub4-mcherry::(KanMX6)

To attach epitope tags to proteins of interest, PCR was used to generate DNA fragments for homologue recombination according to standard procedures (Longtine et al., 1998). Primers and plasmids that were used can be found in the section “Plasmid and primers”. Alternatively, yeast strains were crossed, sporulated (2-3 days at 30°C under agitation) and dissected (Tetrads were treated with sorbitol buffer, 50 mM DTT and 50 μg/mL Zymolase) to achieve the desired combination of tags and deletions within a strain. Tags and deletions were confirmed by colony PCR (Master Mix PCR Hot Start II Phire Green, Thermo Fisher) or microscopy analysis. Yeasts were grown at 30°C with agitation in CSM and exponentially growing cells were harvested for imaging at OD600 0.4.

### Reagents used in the study

Yeast Nitrogen Base w/o AA, Carbohydrate & W/AS (YNB) (USBiological Y2025), Drop-out Mix Complete w/o Yeast Nitrogen Base (DOC) (USBiological D9515), D+ Glucose (USBiological G3050), Nuclease-free water (Ambion-ThermoFisher AM9937), Poly-D-Lysine (Gibco, A3890401); Ammonium persulfate (APS)(17874, ThermoFisher), Tetramethylethylendiamine (TEMED)(ThermoFisher,17919), Formaldehyde 36.5-38% (FA)(SIGMA, F8775) Acrylamide 40% (AA) (SIGMA, A4058), N,N’-methylenbisacrylamide 2% (BIS) (SIGMA, M1533), Sodium Acrylate 97-99% (SA)(SIGMA, 408220), Paraformaldehyde (Electron Microscopy Science, 15700), Zymolyase 100T (USBiological, Z1004), Zymolyase 20T (Roth, 9324.3), Glutaraldehyde (Electron Microscopy Science, cat n°16220). The following primary antibodies were used in this study: YL1/2 anti-a-tubulin (rat) (Abcam ab61610 and gift from Gislene Pereira for *S. pombe*), anti-mcherry (rabbit) (Abcam ab 167453) and anti-centrin (mouse) (Millipore 04-1624, clone 20H5) as well as the indicated secondary antibodies: goat α-mouse 488 mouse (Invitrogen A11029), goat α-mouse 568 (Invitrogen A11004), goat α-rabbit 488 (Invitrogen A11008), goat α-rabbit 568 (Invitrogen A11036), goat α-rat 647 (Invitrogen A21247), goat a-rat 568 (provided by Alba Diz-Munoz).

### Chemicals and solutions

**Potassium phosphate buffer 1M, pH 7.5** was prepared by combining 8.34 mL of 1M K2HPO4 in ddH2O and 1.66 mL of 1M KH2PO4 ddH2O.

**Sorbitol buffer (1.2M)** consisted of 1 mL potassium phosphate buffer 1M, 4 mL 3M sorbitol in ddH2O and 5 mL ddH2O.

**SPO (enhanced Sporulation Medium)** was prepared with 10g/L potassium acetate, 1g/L yeast extract, 0.5g/L D-Glucose and autoclaved

**Sorbitol Buffer** (for dissection) was prepared with 50 mM Tris pH 7.4, 1.2M sorbitol, 5mM EDTA and filtered.

**PFA 16%,** was purchased in 10 mL glass vials, kept at room temperature and before use heated to 37°C for 30 min, this solution was stored at 4°C for up to 1 week. Complete synthetic medium (CSM) contained 20 g/l glucose, 6.706 g/L yeast nitrogen base, 2 g/L drop-out mix complete w/o yeast nitrogen base and 50 mL/L Sörensen buffer.

**CSM (Complete Synthetic Medium)** was prepared 6.7g/L YNB, 2g/L of DOC, 20g/L D-Glucose, 1x Sörensen buffer and sterilized with 0.2um membrane filter.

**CSM/PFA solution** for fixation was prepared fresh right before use, 769 μL CSM, 232 μL of 16% PFA and 110 μL of 1M Potassium phosphate buffer were combined for 1 mL fixative solution.

**Sörensen buffer 20x**, pH 6.2 (0.2M Na2HPO4, 0.8M Kh2PO4) was prepared by dissolving 28.4 g Na2HPO4 and 108.9 g KH2PO4 in 500mL ddH2O and was filtered sterile.

**PEM buffer** consists of 100 mM PIPES, 1 mM EGTA and 1mM MgSO4 and was adjusted to pH 6.9.

**PEMS** was prepared by dissolving 1.2 M sorbitol in PEM buffer.

**PEMBAL** for blocking fission yeast cells consist of 3% (w/v) BSA, 100 mM lysin HCl and 0.1 % (v/v) NaN3.

**Edinburgh Minimal Medium** (EMM) was prepared by mixing 14.7 mM potassium hydrogen phthalate, 15.5 mM Na2HPO4, and 93.5 mM NH4Cl. 2% (w/v) dextrose, salt, minerals, and vitamins were added after autoclavation (Petersen and Russell, 2016)

**Zymolyase** powder was stored at 4°C, aliquots of the solution at −81°C and the running aliquot at −18°C. 50 μL aliquots were prepared by dissolving 25 mg of zymolyase powder in PBS+50% glycerol to yield a final volume of 2.5 mL.

**Anchoring solution** (1.4% formaldehyde / 2% acrylamide (FA / AA) solution in PBS) was prepared fresh right before use. For 1 mL of anchoring solution, precisely 38 μL of FA (36.5-38%), 50 μL of AA (40%) and 912 μL 1X PBS were combined.

**Denaturation buffer** (200 mM SDS, 200 mM NaCl, 50 mM Tris in water, pH 9) was prepared by combining 1.2 g of Tris in 10 mL ddH2O, 114.28 mL SDS 350mM and 8 mL NaCl 5M in 200 mL Of ddH2O. pH was adjusted to 9 by the addition of HCl. 10% TEMED and 10% APS solution was prepared in nuclease free water.

**Monomer solution** (19% Sodium acrylate (SA), 10% Acrylamide (AA), 0.1% Bis-Acrylamide (BIS)) was prepared at least 24 h before use and stored up to 2 weeks at – 18°C. 500 μL of SA, 250 μL of AA, 50 μL BIS and 100 μL 10x PBS were combined for 1 mL of solution.

**Sodium Acrylate** stock solution was prepared as a 38% solution in nuclease-free water by adding 19 g of Sodium Acrylate, little by little, into 31 mL water while stirring. To prepare

**Poly lysine coated coverslips**, coverslips were cleaned by dipping them into 100% ethanol, followed by air-drying. Clean coverslips were put into a humid chamber and incubated with 200 μL (when working with Ø 12 mm coverslips) or 1mL (when working with Ø 24 mm coverslips) of a 100 μg/mL poly-D-lysine solution at 37°C for 45 minutes and washed 3x for 10 min with 200 μL (when working with Ø 12 mm coverslips) or 1 mL (when working with 24 mm coverslips) of ddH2O. Coverslips were stored for up to two weeks at 4°C.

### Yeast culture, cell wall digestion and fixation

*S. cerevisiae* was grown for 36 h in CSM at 30°C with agitation with three dilution cycles or, alternatively was grown on plates for 2 days followed by two rounds of dilution in liquid CSM over a period of 24 h. *S. pombe* was grown in EMM at 32°C for 36 h. For microtubule, SPB and NPC visualization, cells were fixed in 3.7% PFA 0.1M potassium phosphate buffer pH 7.5 (*S. cerevisiae)* or in culture medium (*S. pombe)* for 30-40 min at 21°C with agitation. *S. cerevisiae* were washed with 0.1M phosphate buffer and sorbitol buffer and cell walls were removed by incubation in a mixture of 200 μL sorbitol buffer containing 20 μL of 1M DTT and 1 μL of zymolyase at 1 mg/mL at 30°C until cell walls were removed. In the case of *S. pombe*, cells were washed in PEM buffer to remove residual PFA. Two further washes were performed in PEM containing 1.2 M sorbitol (PEMS) prior to removal the cell walls, which were enzymatically digested with a mixture of 2.5 mg/mL Zymolyase 20T in PEMS at 37°C with agitation for 45 minutes (100 μL/OD600). To check for complete cell wall digestion, equal volumes of Calcofluor white and the digestion mix were combined and imaged at a widefield microscope. Cells were washed 3 times in Sorbitol buffer by centrifugation and resuspended in 200 μL sorbitol buffer. 50 μL of cell suspension was loaded onto a Ø 12 mm poly-lysine-coated coverslip, excess liquid was removed after 10 min. The coverslip was immersed into −20°C ice-cold methanol for 6 min and then immediately into −20°C ice-cold acetone for 30 sec and allowed to dry.

### U-ExM protocol

Samples were fixed and cell walls were digested as in IF analysis. Poly-lysine-coated coverslips with fixed spheroplasts were incubated in protein crosslinking prevention solution (2% AA / 1.4% FA in PBS) for 3 to 5h at 37°C. Gelation was performed on ice, APS and TEMED were added to the monomer solution (19% (wt/wt) SA, 10% (wt/wt) AA, 0.1% (wt/wt) BIS in 1X PBS) to a final concentration of 0.5% each. 35 μL of this solution was covered with the prepared coverslip in a pre-cooled humid chamber. The humid chamber containing the coverslips was incubated 5 min on ice and then for 1 h at 37°C. Sample coverslips were incubated in denaturation buffer with agitation for 15 min at RT to facilitate gel detachment from the coverslips. Gels were then transferred to Eppendorf tubes filled with fresh denaturation buffer and incubated for 1 h 30 min at 95°C without agitation. After denaturation, gels were expanded with three subsequent baths of ddH2O for 30 min at RT. After full expansion of the gel, the diameter of the gel was measured and proceed for immunostaining (see section below).

### Immunofluorence staining

For pre-expansion pan labelling*, S. cerevisiae* cells were incubated in PBS-BSA 1% for 10 min, washed three times with PBS and stained with NHS-ester diluted at 2μg/mL in PBS for 1 h 30 min at RT in the dark. The coverslip was washed three times with PBS. The coverslips were mounted onto a glass slide using a glycerol based mounting medium containing DAPI. *S. pombe* incubated with NHS-ester diluted at 2μg/mL in PBS over night at 4°C. The coverslips were washed three times with PEM and Hoechst 33342 was added at 0.25 μg/mL in PBS for 5 minutes before mounting coverslips in ProLong Diamond Antifade mountant.

For post-expansion staining, expanded gels were incubated in PBS for 2x 15 min at RT. Gels were stained in PBS-BSA 2% containing primary antibody for 2 h 30 min at 37°C with agitation and the gel was washed 3x with PBS Tween 0.1% for 10 min at RT with agitation. Gels were then incubated with PBS-BSA 2% containing secondary antibody for 2 h 30 min at 37°C with agitation in the dark and the gel was washed 3x with PBS Tween 0.1% for 10 min at RT with agitation in the dark. The gel is incubated in water for 2x 30 min at RT and is than left to fully expand overnight in fresh water in the dark before imaging. For *S. pombe* IF, antibody dilutions were prepared in PEM buffer with 2 % BSA and 0.2 % Tween-20. The gels were incubated on a rotating wheel in primary antibodies over night at 4°C prior to washing them in PEM buffer with 0.2 % Tween-20) three times. Gels were incubated with PEM buffer with 2 % BSA and 0.2 % Tween-20 containing secondary antibody for 2 h 30 min at 37°C with agitation. The samples were subsequently washed three times with PBS-T before expanding them as described above. For pan labeling, gels were incubated in 2μg/mL NHS-ester in PBS over night at 4°C without agitation for *S. pombe* and in 10 μg/mL NHS-ester in PBS 1h30 at RT with agitation for *S. cerevisiae.* Gels were next washed 3-4 times during 10min with PBS tween prior to imaging. Note that while NHS-ester incubation after IF is possible, this may lead to increased signals colocalizing with antibody signals.

### Sample mounting and imaging

For gel imaging, a piece of approximately 1×1 cm^2^ was cut from the center of the gel and the backside of the gel, which does not contain cells, was slightly dried. The gel was then attached to a 24 mm poly-lysine-coated coverslip or Ibidi chamber with the front, cell-containing side of the gel touching the glass. The coverslip was mounted into a metal holder, which can be attached to the microscope and a drop of water was added onto the gel before imaging to prevent drying of the gel. Confocal and widefield imaging were performed as previously published (Gambarotto et al., 2019; Gambarotto et al., 2021). For *S. cerevisiae*, confocal imaging was performed using a Leica TCS SP8 microscope with a 63x oil-immersion objective with 1.4-NA (numerical aperture) in the lightening mode at max resolution, generating deconvolved images. Water was considered as mounting medium and adaptive strategy was chosen. The step size for z-stack acquisitions was 0.12 μm, with a pixel size of 35 nm. Widefield imaging was performed using a Leica DM18 microscope with a 63x oil immersion objective with 1.4-NA with the Thunder “Small volume computational clearing” mode to generate deconvolved images. For gel imaging, water was considered as mounting medium while Vectashield was used for regular immunofluorescence. Pictures were acquired with a z-stack size of 0.21 μm using a pixel size of 100 nm. For *S. pombe*, an Olympus IXplore SpinSR spinning disk confocal with a 40x NA 0.95 air objective for overview images and a 63x oil-immersion objective (NA 1.42) was used for width determination. Z-stacks were aquired at a step size of 0.3 μm. For NPC counting, gels were imaged at a Zeiss LSM980 AiryFast confocal microscope using a 63x oil immersion objective (NA 1.4) at a step size of 0.15 μm. Widefield images were taken at a Zeiss CellObserver with an 63x objective (NA 1.4) at 0.5 μm z-slices.

### Quantification and statistical analysis

#### Width/diameter measurements

The distance between the half maximum intensity of the first and last signals along a line across the cell was determined using the plot profile tool of Fiji. For *S. cerevisiae*, two perpendicular measurements were applied and averaged for every cell, due to oval shape of yeast. For *S. pombe* width determination, the diameter of the cell was measured perpendicular to the length across the nucleus.

#### Dimensions of the SPB in S. cerevisiae

The distance of the outer and inner plaques of the SPB was measured as the x-axis distance between the maximum intensities of both signals corresponding to the two plaques. Sfi1 and Cdc31 signal length was determined as for the cellular diameters, always starting the measurement from the end of the signal that overlaps with the spindle and thus presumably corresponds to the side of the bridge that is closer to the old SPB.

#### NPCs quantification in S. pombe

NPCs were quantified using the 3D objects counter FIJI plugin (Bolte and Cordelieres, 2006) to determine centroids and the number of objects. Manual thresholds were set to adjust for variations in staining intensities and objects were manually cleaned to remove unspecific signals, using the pan labelled nuclear envelope as a reference. Cell cycle stages were defined using the adjusted cell length, taking the expansion factor of 4.2-fold as a basis for calculations, as previously described (Varberg et al., 2021). In brief, mononucleated cells below 9.5 μm were classified as early G2 and those between 9.5 μm and 11 μm as mid G2. Cells longer than 11 μm were defined as late G2/M when mononucleated, while in binucleated cells the state of the septum was considered to differentiate late M (no full separation) from G1/S cells (septum closed).

### Data availability

All data are available in this manuscript._Further information and request for resources can be addressed to Paul Guichard and Virginie Hamel. *S. cerevisiae* strain requests should be addressed to Robbie Loewith and *S. pombe* strain requests to Gautam Dey.

